# Novel Significant Stage-Specific Differentially Expressed Genes in Liver Hepatocellular Carcinoma

**DOI:** 10.1101/342204

**Authors:** Arjun Sarathi, Ashok Palaniappan

**Affiliations:** Depts. of Bioengineering, School of Chemical and BioTechnology, SASTRA deemed University, Thanjavur, Tamil Nadu 613401. INDIA; Depts. of Bioinformatics, School of Chemical and BioTechnology, SASTRA deemed University, Thanjavur, Tamil Nadu 613401. INDIA

**Keywords:** LIHC transcriptomics, HCC stages, stage-specific biomarkers, differentially expressed genes, pairwise contrasts, significance analysis, linear modelling, tumorigenesis, cancer progression, metastasis

## Abstract

Liver cancer is among the top deadly cancers worldwide with a very poor prognosis, and the liver is a particularly vulnerable site for metastasis of other cancers. In this study, we developed a novel computational framework for the stage-specific analysis of hepatocellular carcinoma initiation and progression. Using publicly available clinical and RNA-Seq data of cancer samples and controls, we annotated the gene expression matrix with sample stages. We performed a linear modelling analysis of gene expression across all stages and found significant genome-wide changes in gene expression in cancer samples relative to control. Using a contrast against the control, we were able to identify differentially expressed genes (log fold change >2) that were significant at an adjusted p-value < 10E-3. In order to identify genes that were specific to each stage without confounding differential expression in other stages, we developed a full set of pairwise stage contrasts and enforced a p-value threshold (<0.05) for each such contrast. Genes were specific for a stage if they passed all the significance filters for that stage. Our analysis yielded two stage-I specific genes (CA9, WNT7B), two stage-II specific genes (APOBEC3B, FAM186A), ten stage-III specific genes including DLG5, PARI and GNMT, and ten stage-IV specific genes including GABRD, PGAM2 and PECAM1. Of these, only APOBEC3B is an established cancer driver gene. DLG5 was found to be tumor-promoting contrary to the cancer literature on this gene. Further, GABRD, well studied in literature on other cancers, emerged as a stage-IV specific gene. Our findings could be validated using multiple sources of omics data as well as experimentally. The biomarkers identified herein could potentially underpin diagnosis as well as pinpoint drug targets.

## INTRODUCTION

Liver cancer is the second most deadly cancer in terms of mortality rate, with a very poor prognosis (Yang et al 2010). It accounted for 9.1% of all cancer deaths, and 83% of the annual new estimated 782,000 liver cancer cases worldwide occur in developing countries (Ferlay et al., 2015). Liver cancer showed the greatest increase in mortality in the last decade for both males (53%) and females (59%) (Cancer Research UK, 2018). Liver hepatocellular carcinoma (LIHC; HCC) is the most common type of liver cancer. 78% of all reported cases of LIHC were due to viral infections (53% Hepatitis B virus and 25% Hepatitis C virus) (Perz et al., 2006). There are several non-viral causes of LIHC as well, mainly aflatoxins and alcohol (Chuang et al., 2009). As shown in Fig. 1, all the factors converge to a common mechanism of genetic alterations that lead to the acquisition of cancer hallmarks (Hanahan and Weinberg, 2011) and the eventual emergence of a cancer cell (Farazi et al., 2006). Genetic alterations constitute the heart of the problem, and studying changes due to these genetic alterations is paramount to understand LIHC. Early gene expression studies using EST data detected differential expression in cancer tissue compared to non-cancerous liver and proposed the existence of genetic aberrations and changes in transcriptional regulation in LIHC (Xu et al., 2001). The Cancer Genome Atlas (TCGA) research network (2017) have subtyped and identified many potential targets for LIHC based on a comprehensive multi-omics analysis. An independent analysis of TCGA RNA-Seq data encompassing 12 cancer tissues has uncovered liver cancer-specific genes (Peng et al., 2015). Zhang et al. (2015) have performed mutation analysis of LIHC, and Yang et al. (2017) combined TCGA expression data and natural language processing techniques to identify cancer-specific markers.

**Figure 1.**
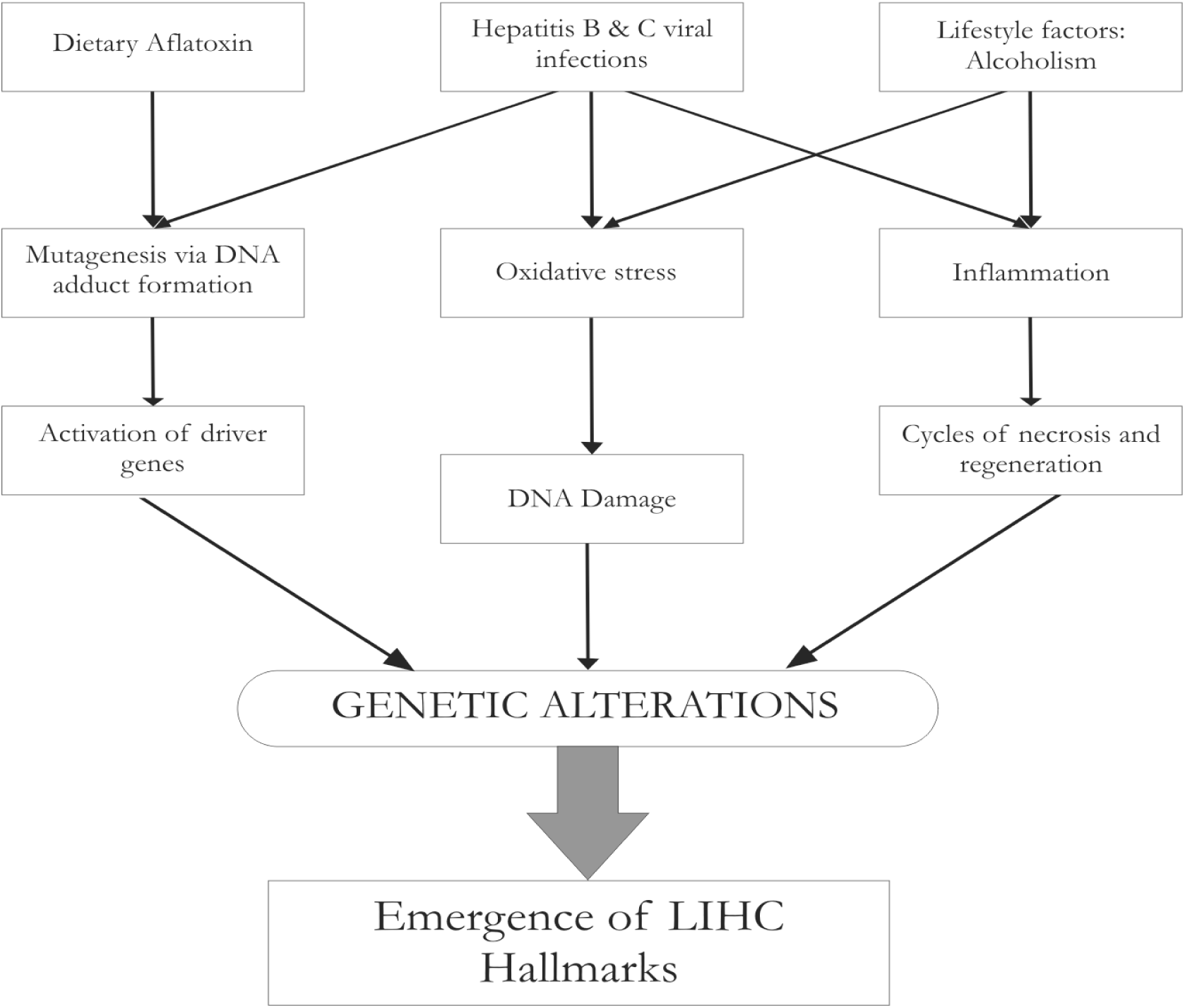
Major causative pathways of hepatocarcinogenesis. All pathways converge to progressive genomic alterations, leading a normal cell to acquire the hallmarks of cancer.

The burden of disease and mortality rate are both inversely correlated with the cancer stage. The response rate to therapy is also inversely correlated with stage. To the best of our knowledge, there are no reported research in the literature that have dissected the stage-specific features of LIHC. The cancer staging system is based on gross features of cancer anatomical penetration, and one such standard is the American Joint Committee on Cancer (AJCC) Tumor-Node-Metastasis (TNM) staging (Amin et al., 2017). It is reasonable to hypothesize that the stage-specific gross changes are associated with signature molecular events, and try to probe such molecular bases of stage-wise progression of cancer. We had earlier published on stage-specific “hub driver” genes in colorectal cancer (Palaniappan et al., 2016). A stage-focussed analysis of colorectal cancer transcriptome data yielded negative results vis-a-vis the AJCC staging system (Huo et al., 2017).

## METHODS

### DATA PREPROCESSING

Normalized and log2-transformed Illumina HiSeq RNA-Seq gene expression data processed by the RSEM pipeline (Li and Dewey, 2011) were obtained from TCGA via the firebrowse.org portal (Broad Institute TCGA Genome Data Analysis Center, 2016). The patient barcode (uuid) of each sample encoded in the variable called ‘Hybridization REF’ was parsed and used to annotate the controls and cancer samples (Fig. 2). To annotate the stage information of the cancer samples, we obtained the clinical information dataset for LIHC from firebrowse.org (LIHC.Merge_Clinical.Level_1.2016012800.0.0.tar.gz) and merged the clinical data with the expression data by matching the “Hybridization REF” in the expression data with the aliquot barcode identifier in the clinical data. The stage information of each patient was encoded in the clinical variable “pathologic stage”. The substages (A,B,C) were collapsed into the parent stage, resulting in four stages of interest (I, II, III, IV). We retained a handful of other clinical variables pertaining to demographic features, namely age, sex, height, weight, and vital status. With this merged dataset, we filtered out genes that showed little change in expression across all samples (defined as σ < 1). Finally, we removed cancer samples from our analysis that were missing stage annotation (value ‘NA’ in the “pathologic stage”). The data pre-processing was done using R (www.r-project.org).

**Figure 2.**
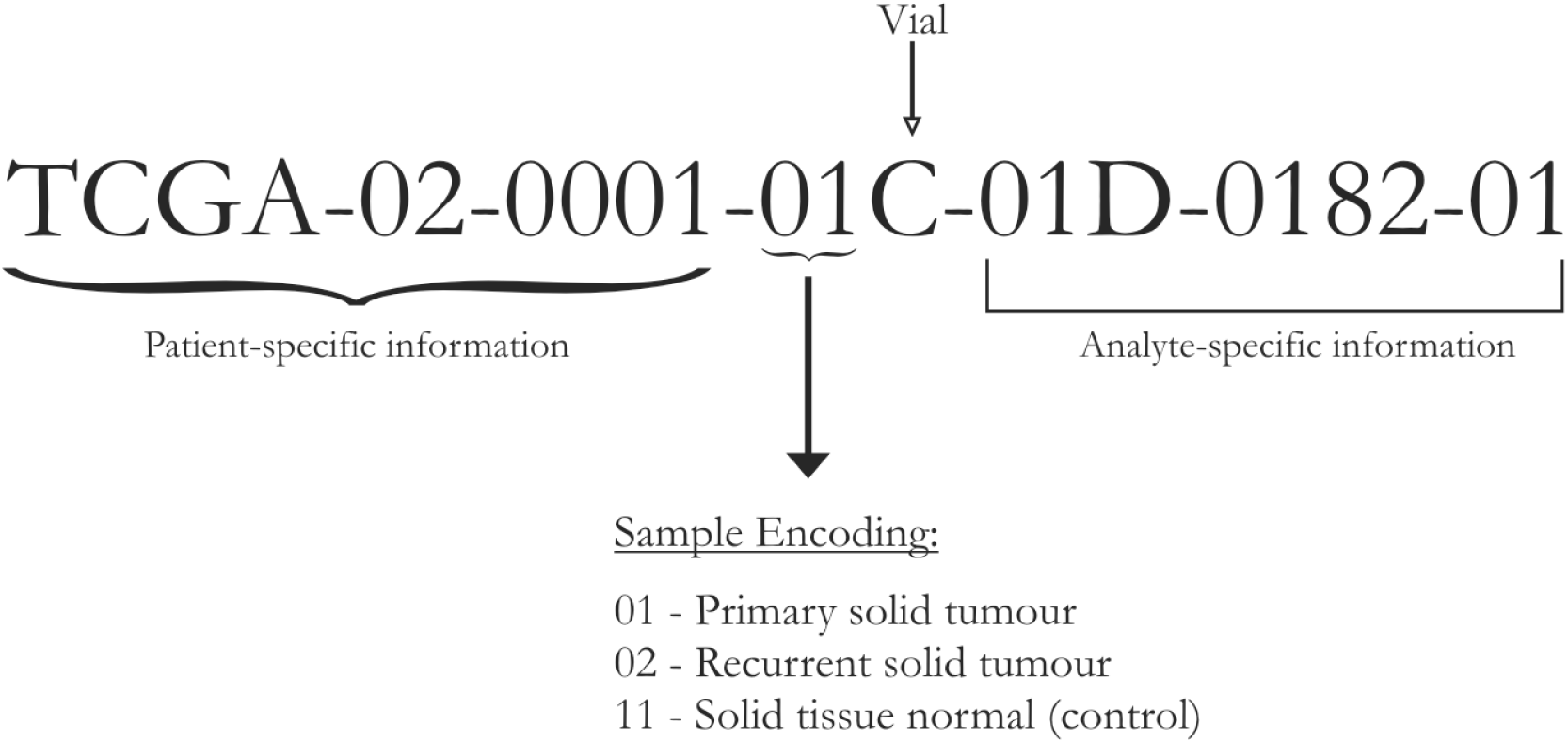
TCGA ‘Hybridization REF’ Barcode. The first 10 characters constitute an anonymized unique patient identifier and the following two characters denote whether the sample is tumor or normal.

### LINEAR MODELLING

Linear modelling of expression across cancer stages relative to the baseline expression (i.e, in normal tissue controls) was performed for each gene using the R *limma* package (Ritchie et al., 2015). The following linear model was fit for each gene’s expression based on the design matrix shown in Fig. 3A:

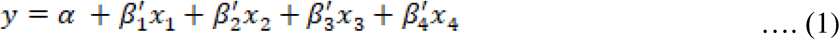

where the independent variables are indicator variables of the sample’s stage, the intercept α is the baseline expression estimated from the controls, and β_i_ are the estimated stagewise log fold-change (lfc) coefficients relative to controls. The linear model was subjected to empirical Bayes adjustment to obtain moderated t-statistics (McCarthy and Smyth, 2009). To account for multiple hypothesis testing and the false discovery rate, the p-values of the F-statistic of the linear fit were adjusted using the method od Hochberg and Benjamini (1990). The linear trend across cancer stages for the top significant genes were visualized using boxplots to ascertain the regulation status of the gene relative to the control.

**Figure 3.**
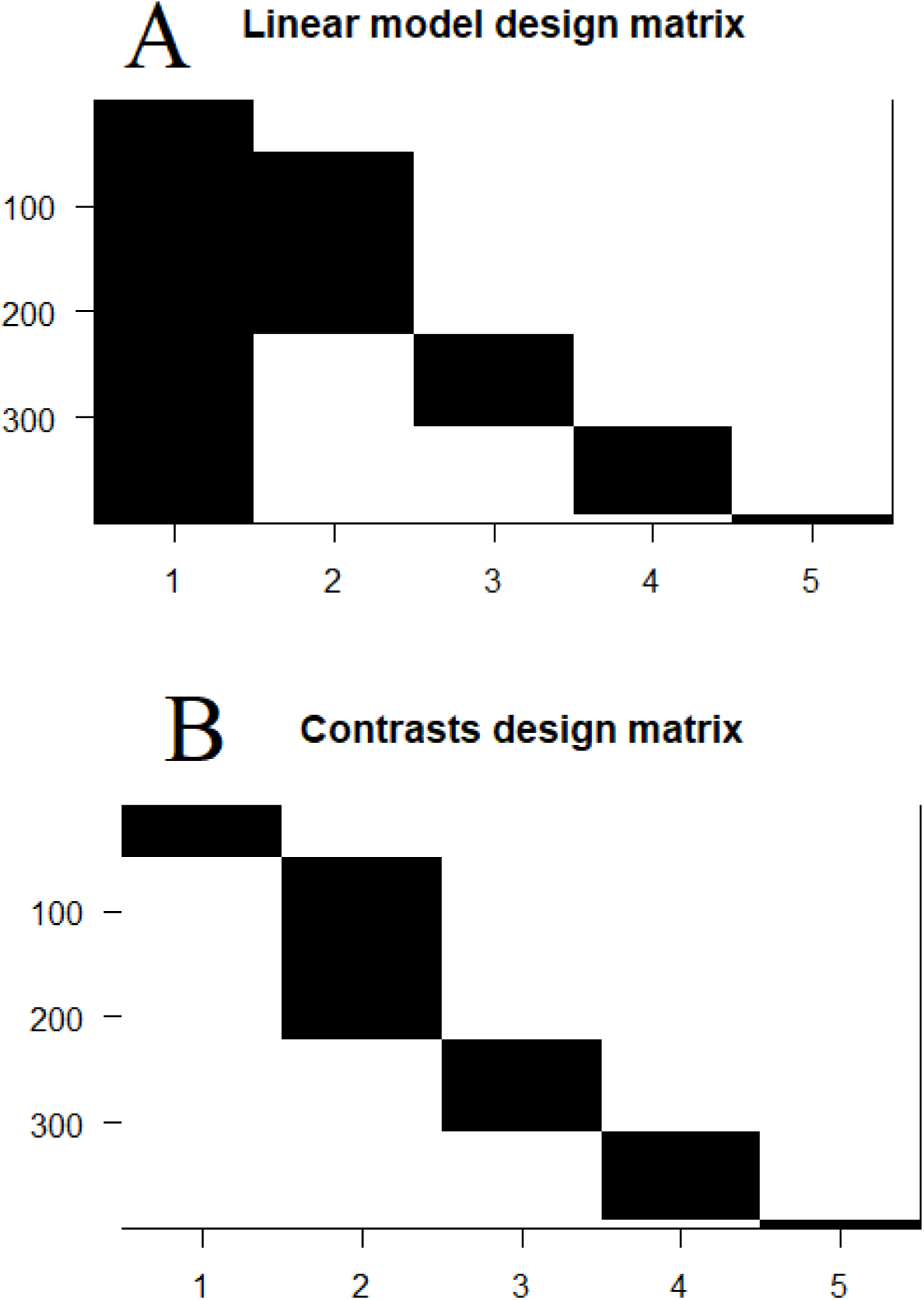
Design matrices. **A,** In the linear modeling, the control samples served as the baseline expression (intercept) of each gene against which the stage-specific expression was estimated. **B,** the design matrix for the contrasts analysis.

### PAIRWISE CONTRASTS

To perform contrasts, a slightly modified design matrix shown in Fig. 3B was used, which would give rise to the following linear model of expression for each gene:

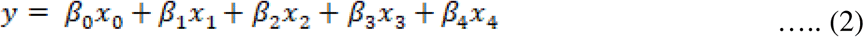

where the controls themselves are one of the indicator variables, and the βi are all coefficients estimated only from the corresponding samples. Our first contrast of interest, between each stage and the control, was achieved using the contrast matrix shown in Table 1. Four contrasts were obtained, one for each stage vs control. A threshold of |lfc| > 2 was applied to each such contrast to identify differentially expressed genes (with respect to the control). We used the absolute value of the lfc, since driver genes could be either upregulated or downregulated. Genes could be differentially expressed in any combination of the stages or no stage at all. To analyze the pattern of differential expression (with respect to the control), we constructed a four-bit binary string for each gene, where each bit signified whether the gene was differentially expressed in the corresponding stage. For example, the string ‘1100’ indicates that the gene was differentially expressed in the first and second stages. There are 2^4^ =16 possible outcomes of the four-bit string for a given gene corresponding to the combination of stages in which it is differentially expressed. This is illustrated in set-theoretic terms in Fig. 4. In our first elimination, we removed genes whose |lfc| < 2 for all stages. For each remaining gene, we identified the stage that showed the highest |lfc| and assigned the gene as specific to that stage for the rest of our analysis.

**Figure 4.**
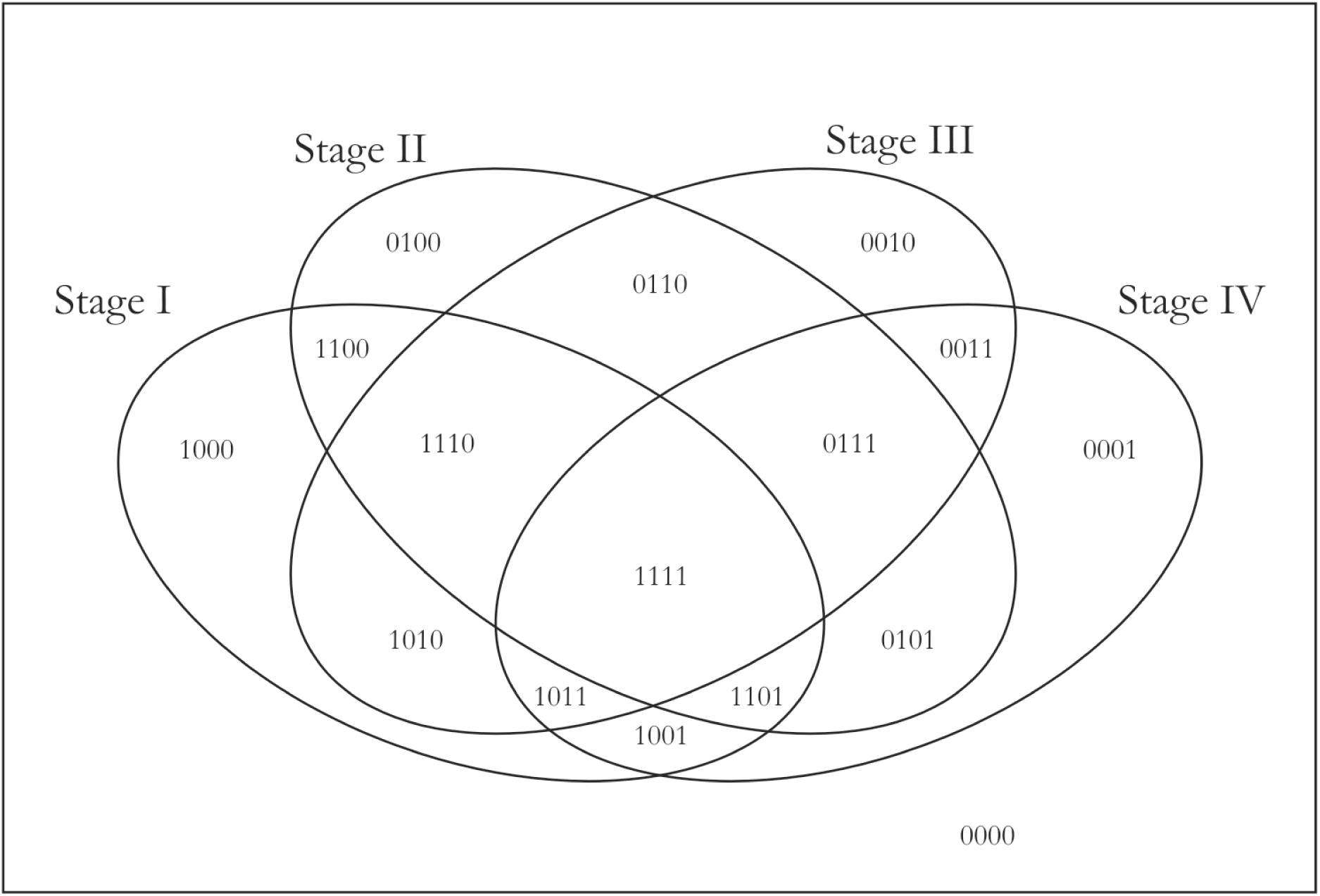
A Venn representation of the pairwise stages contrasts. A gene could be differentially expressed in any combination of the four stages and this could be represented by a 4-bit string, one bit for each stage. For e.g, ‘1111’ at the overlap of all four stages would be assigned to genes that are differentially expressed in all four stages.

**Table 1.** Contrast matrix with control. Each stage (indicated by ‘1’) is contrasted against the control (indicated by ‘-1’) in turn.

|  | Stage1-control | Stage2-control | Stage3-control | Stage4-control |
| --- | --- | --- | --- | --- |
| Control | -1 | -1 | -1 | -1 |
| Stage1 | 1 | 0 | 0 | 0 |
| Stage2 | 0 | 1 | 0 | 0 |
| Stage3 | 0 | 0 | 1 | 0 |
| Stage4 | 0 | 0 | 0 | 1 |

### SIGNIFICANCE ANALYSIS

We applied a four-pronged criteria to establish the significance of the stage-specific differentially expressed genes.

(i) Adj. p-value of the contrast with respect to the control < 0.001

(ii)-(iv) P-value of the contrast with respect to other stages < 0.05

To obtain the above p-values (ii) - (iv), we used the contrast matrix shown in Table 2, which was then used an an argument to the `contrastsFit` function in *limma*.

**Table 2.** Contrast matrix for inter-stage contrasts. There are six possible pairwise contrasts between the stages that are essential to identifying stage-specific genes.

|  | Stage2-Stage1 | Stage3-Stage1 | Stage3-Stage2 | Stage4-Stage1 | Stage4-Stage2 | Stage4-Stage3 |
| --- | --- | --- | --- | --- | --- | --- |
| Control | 0 | 0 | 0 | 0 | 0 | 0 |
| Stage1 | -1 | -1 | 0 | -1 | 0 | 0 |
| Stage2 | 1 | 0 | -1 | 0 | -1 | 0 |
| Stage3 | 0 | 1 | 1 | 0 | 0 | -1 |
| Stage4 | 0 | 0 | 0 | 1 | 1 | 1 |

### FURTHER ANALYSES

Principal component analysis (PCA) were performed using `prcomp` in R. To choose 100 random genes, we used the `rand` function. Gene set enrichment analysis were performed on KEGG (www.kegg.ac.jp) and Gene Ontology (Ashburner et al., 2000) using `kegga` and `goana` in *limma*, respectively. In order to visualize outlier genes that are significant with a large effect size, volcano plots could be obtained by plotting the -log10 transformed p-value vs. the log fold-change of gene expression. Heat maps of significant stage-specific differentially expressed genes were visualized using `heatmap` and clustered using `hclust`. Novelty of the identified stage-specific genes was ascertained by screening against the Cancer Gene Census v84 (Futreal et al., 2004).

## RESULTS

The TCGA expression data consisted of expression values of 20,532 genes in 423 samples. After the completion of data pre-processing, we obtained a final dataset of expression data for 18,590 genes across 399 samples annoatated with the corresponding sample stage (available in Supplementary File S1). The stagewise distribution of TCGA samples along with the corresponding AJCC staging is shown in Table 3. A statistical summary of demographic details including age, sex, height, weight, and vital status is shown in Table 4. The body mass index (BMI) distribution was derived from patient clinical data that had both height and weight (i.e, neither was ‘NA’). The average age of onset of LIHC was around 60 years, and the average BMI was about 26, indicating a possible link with ageing and obesity.

**Table 3.**
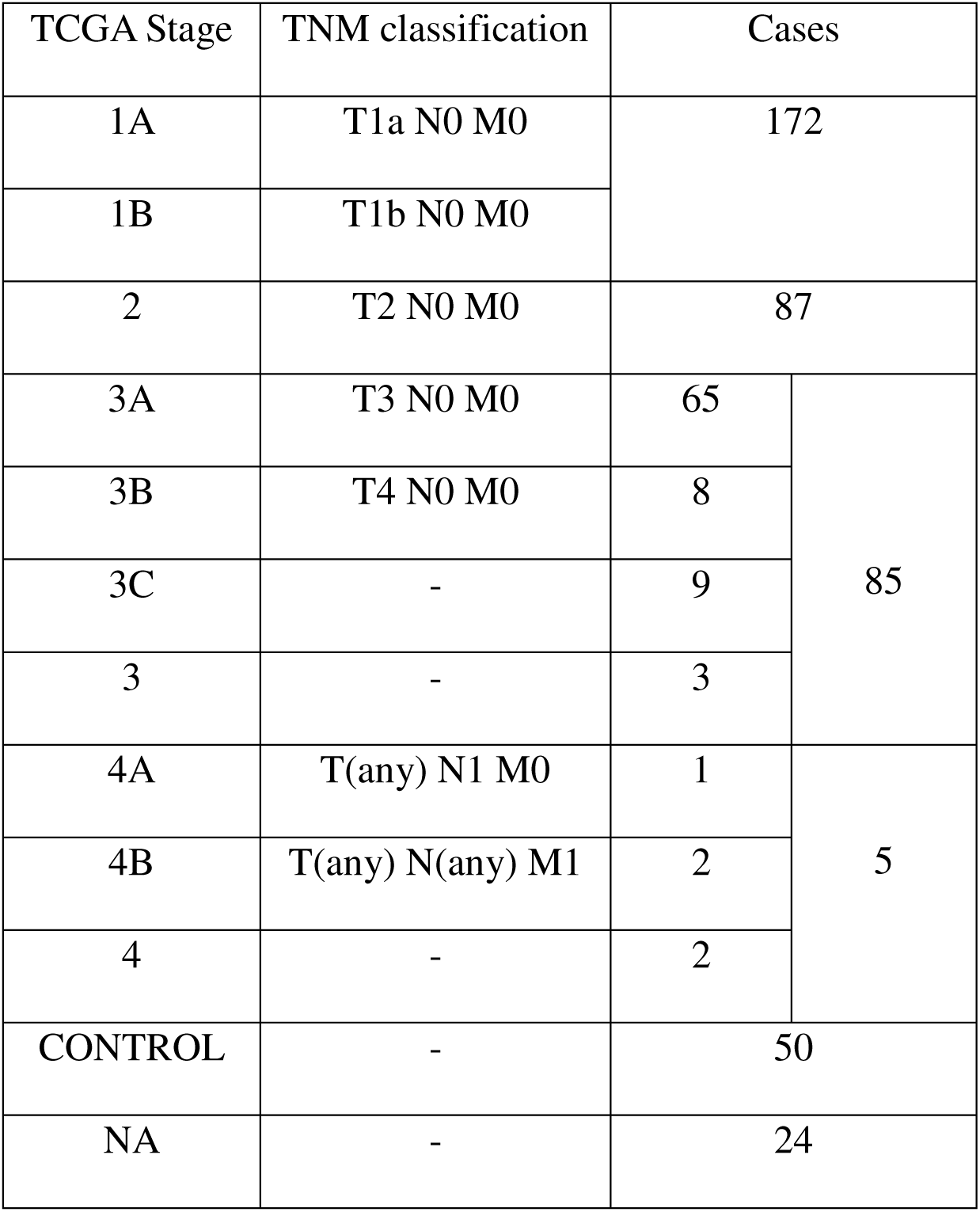
AJCC Cancer staging. The correspondence between the AJCC staging and the TCGA staging for LIHC is noted, along with the number of LIHC cases in each stage in the TCGA dataset. Control indicates the number of normal tissue control samples, and NA denotes cases where the stage information is unavailable.

**Table 4.** Summary of key demographic features of the dataset. For continuous variables (age, height, weight and BMI), the mean ± standard deviation is given. BMI is calculated only for patients with both height and weight data.

| Characteristic |  | Control | Stage1 | Stage2 | Stage3 | Stage4 | NA | Overall |
| --- | --- | --- | --- | --- | --- | --- | --- | --- |
| Number of samples |  | 50 | 172 | 87 | 85 | 5 | 24 | 423 |
| Age (Years) |  | 61.7±16.1 | 60.6±12.2 | 59.0±13.3 | 56.2±14.8 | 42.8±20.7 | 68.1±10.7 | 59.7±13.2 |
| Height (cm) |  | 170.6±9.5 | 166.5±12.3 | 167.9±8.3 | 169.0±8.9 | 162.3±4.9 | 166.2±11.1 | 167.7±10.6 |
| Weight (Kg) |  | 76.1±22.1 | 73.2±19.8 | 73.3±18.9 | 69.9±18.8 | 72.3±21.5 | 79.7±19.8 | 73.2±19.7 |
| BMI |  | 26.2±7.8 | 26.7±10.2 | 26.0±5.8 | 24.3±6.0 | 27.7±9.3 | 29.1±7.6 | 26.1±8.4 |
| Sex | Male | 28 | 122 | 60 | 55 | 1 | 14 | 280 |
|  | Female | 22 | 50 | 27 | 30 | 4 | 10 | 143 |
| Vital Status | Alive | 20 | 134 | 71 | 65 | 2 | 12 | 304 |
|  | Dead | 30 | 38 | 16 | 20 | 3 | 12 | 119 |

The dataset was processed through `voom` in *limma* to prepare for linear modelling (Law et al., 2014). At a p-value cutoff of 10E-5, 9618 genes were significant in the linear modelling, implying a strong linear trend in their expression across cancer stages. This was not entirely surprising since one of the hallmarks of cancer phenotype is genome-wide instability (Hanahan and Weinberg, 2011). The linear modelling highlighted top ranked genes, some upregulated in LIHC (GABRD, PLVAP, CDH13) and some downregulated (CLEC4M, CLEC1B, CLEC4G). The lfc for each stage with respect to control of top ten genes (ranked by adjusted p-value) are shown in Table 5, along with their inferred regulation status. Boxplots of the expression of the top 9 genes (Fig 5) indicated a progressive net increase in expression across cancer stages relative to control for up-regulated genes, while depressed expression across cancer stages relative to control was indicative of downregulated genes. (Boxplots of all other genes in the top 200 are provided in the Supplementary Fig. S1) It is worthwhile to note that a given gene might have maximal differential expression in any stage (not necessarily stage 4), and the linear trend does not suggest the order of expression across stages (Fig. 6).

**Figure 5.**
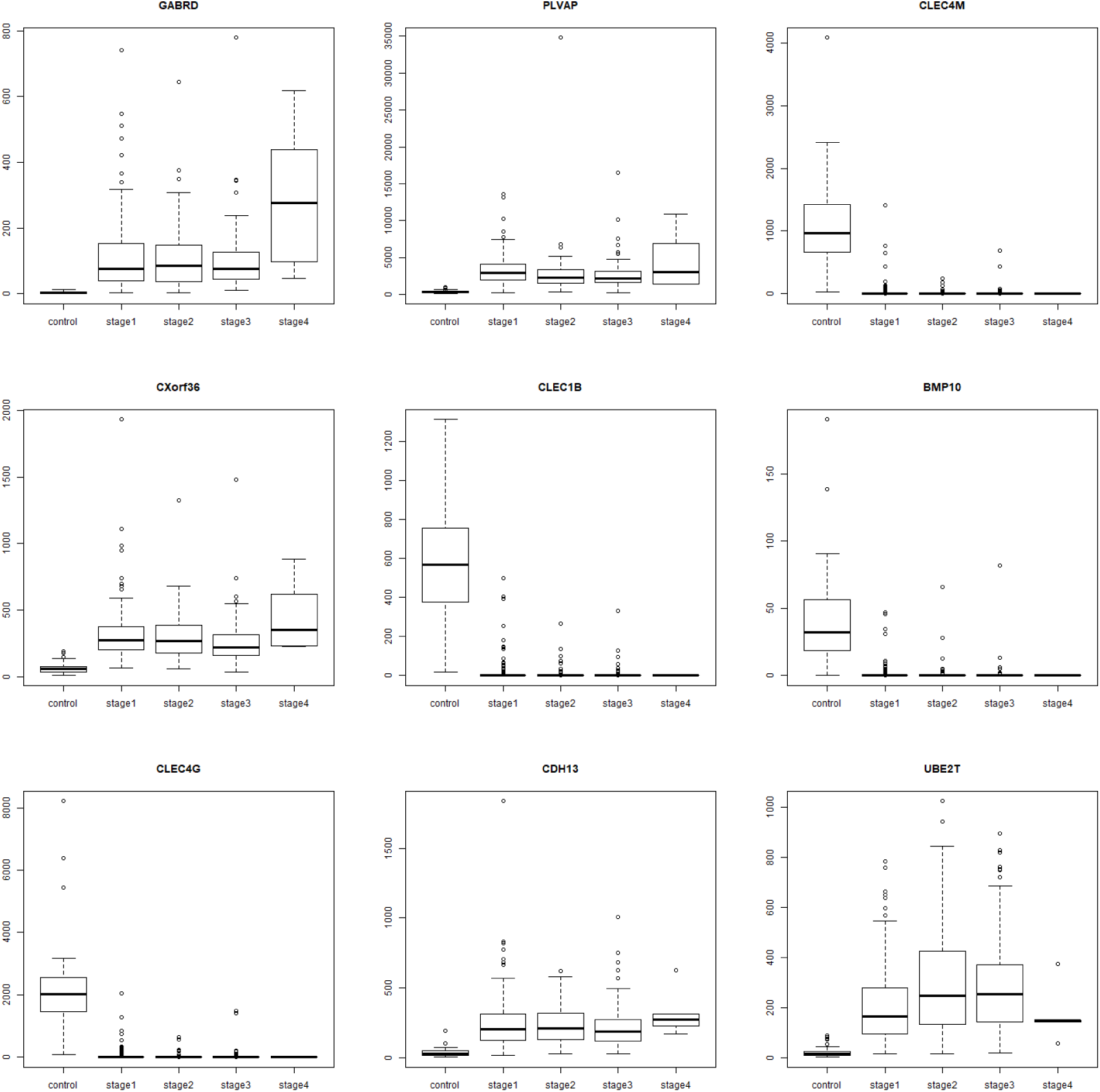
Boxplots of top 9 linear model genes. For each gene, notice that the trend in expression could be overexpression or downregulations relative to the control. For e.g, GABRD, PLVAP, CXorf36, CDH13 and UBE2T are overexpressed, while CLEC4M, CLEC1B, BMP10, and CLEC4G are downregulated. It could be seen that a linear trend does not imply maximal |lfc| in stage 4, as illustrated most clearly in the case of UBE2T.

**Table 5.** Top 10 genes of the linear model. The log-fold change expression of the gene in each stage relative to the controls are given, followed by p-value adjusted for the false discovery rate, and the regulation status of the gene in the cancer stages with respect to the control.

| Genes | Stage I<br>lfc ($\beta_1$) | Stage II<br>lfc ($\beta_2$) | Stage III<br>lfc ($\beta_3$) | Stage IV<br>lfc ($\beta_4$) | Adj. p<br>value | Regulation status |
| --- | --- | --- | --- | --- | --- | --- |
| GABRD | 5.08 | 5.11 | 5.24 | 6.55 | 5.529e-78 | Up-regulated |
| PLVAP | 3.51 | 3.24 | 3.24 | 3.79 | 7.498e-75 | Up-regulated |
| CLEC4M | -8.32 | -8.67 | -8.48 | -9.24 | 6.058e-74 | Down-regulated |
| CXORF36 | 2.91 | 2.86 | 2.76 | 3.44 | 5.376e-73 | Up-regulated |
| CLEC1B | -7.85 | -8.46 | -8.05 | -9.44 | 6.292e-71 | Down-regulated |
| BMP10 | -4.66 | -4.75 | -4.67 | -5.25 | 1.447e-66 | Down-regulated |
| CLEC4G | -7.75 | -8.23 | -7.95 | -8.75 | 2.437e-66 | Down-regulated |
| CDH13 | 3.30 | 3.34 | 3.32 | 3.86 | 3.454e-66 | Up-regulated |
| UBE2T | 3.85 | 4.50 | 4.47 | 3.76 | 2.544e-65 | Up-regulated |
| SLC26A6 | 3.10 | 3.39 | 3.34 | 3.07 | 7.438e-65 | Up-regulated |

**Figure 6.**
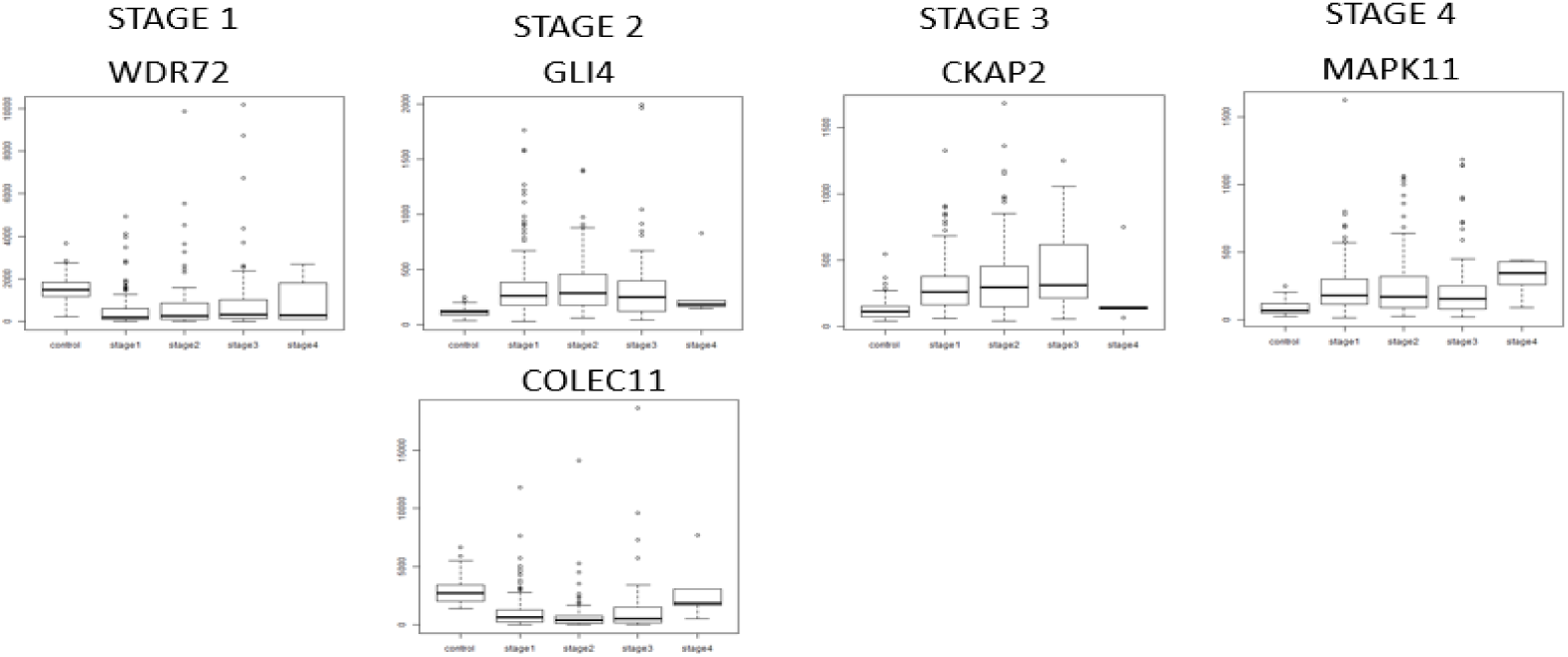
Boxplots illustrating stage-specificity of differentially expressed genes. Extremal expression in a stage could be either maximal expression or minimal expression relative to the control and all other stages, and could be termed maximal differential expression. Here we show genes with maximal differential expression in stage-I (WDR72; minimum expression), stage-II (GLI4, maximum expression; COLEC11, minimum expression), stage-III (CKAP2; maximum expression), and stage-IV (MAPK11; maximum expression).

A PCA of the top 100 genes from the linear model was visualized using the top two principal components (Fig. 7A). A clear separation of the controls and the cancer samples could be seen, suggesting the extent of differential expression of these genes in cancer samples. Hence linear modelling yields cancer-specific genes versus normal controls, and the results for the all the genes, including the top 100, are provided in order in Supplementary File S2. For comparison, a PCA plot of 100 randomly sampled genes (Fig. 7B) failed to show any separation of the cancer and control samples.

**Figure 7.**
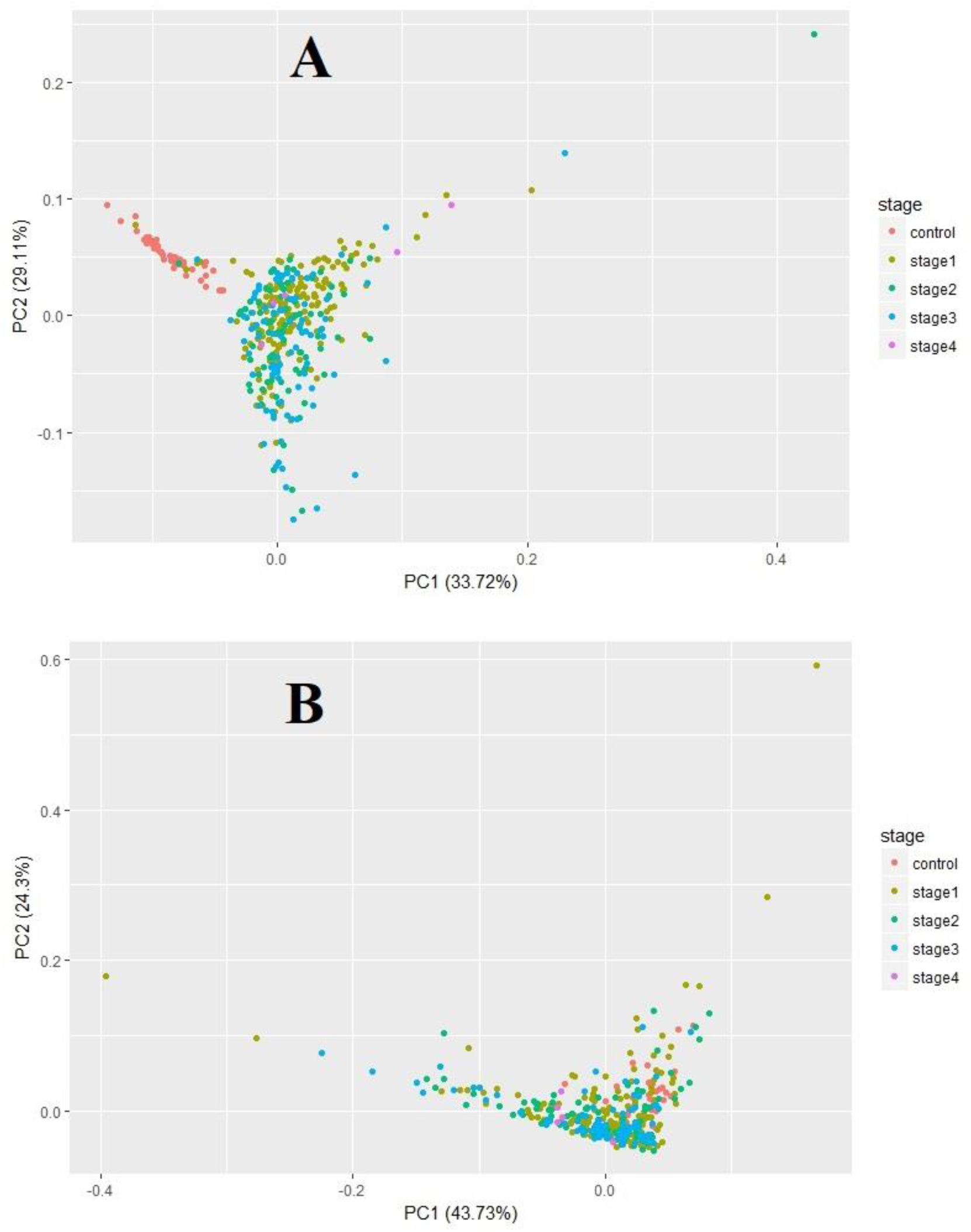
Principal components analysis of cancer vs control. **A,** The first two principal components of the top 100 genes from linear modeling are plotted. It could be seen that control samples (red) clustered independent of the cancer samples (colored by stage). **B,** The same analysis repeated with 100 random genes failed to effect a clustering of the control samples relative to the cancer samples.

The results from the linear modelling were in contrast with those obtained by Huo *et al*. (2017) and were most likely driven by the inclusion of 51 controls in our study. These positive results provided the impetus to pursue stage-driven analysis. Given the conventional AJCC staging, gene expression differences would play a major role in driving the cancer progression. To identify the stage-specific differentially expressed genes, we applied the first contrast matrix (Table 2) and constructed the four-bit stage string of each gene. Based on the stage strings, we binned all the genes, and the string-specific gene lists corresponding to all the partitions in the Venn diagram (Fig. 4) is made available in Supplementary File S3. The size of each such partition is illustrated in Fig. 8. We eliminated the 16,135 genes corresponding to the stage string ‘0000’ ( |lfc|<2 in all stages). To establish the significance of the remaining genes, we applied the second contrast (Table 3) and passed each gene through the four filter criteria. The gradual reduction in candidate stage-specific genes as each criterion was applied, is shown in Table 6. Only genes that passed all criteria were retained as significant stage-specific differentially expressed genes. We obtained 2 stage-I specific, 2 stage-II specific, 10 stage-III specific and 35 stage-IV specific genes (Table 7). Fig. 9 shows the volcano plot of these 49 stage-specific genes.

**Figure 8.**
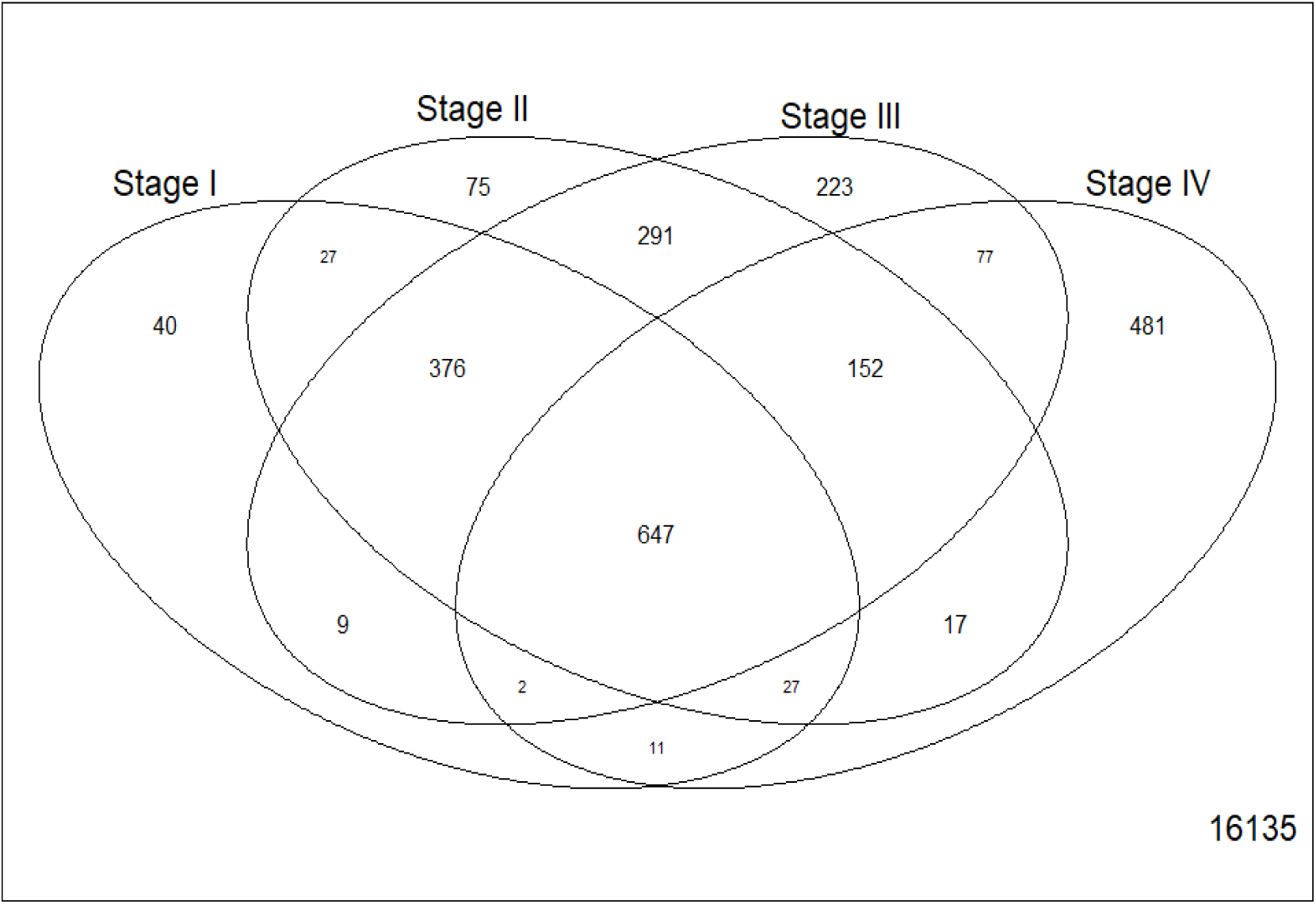
Venn illustration of the size of each 4-bit string. The number of genes having each pattern of differential expression are shown.

**Figure 9.**
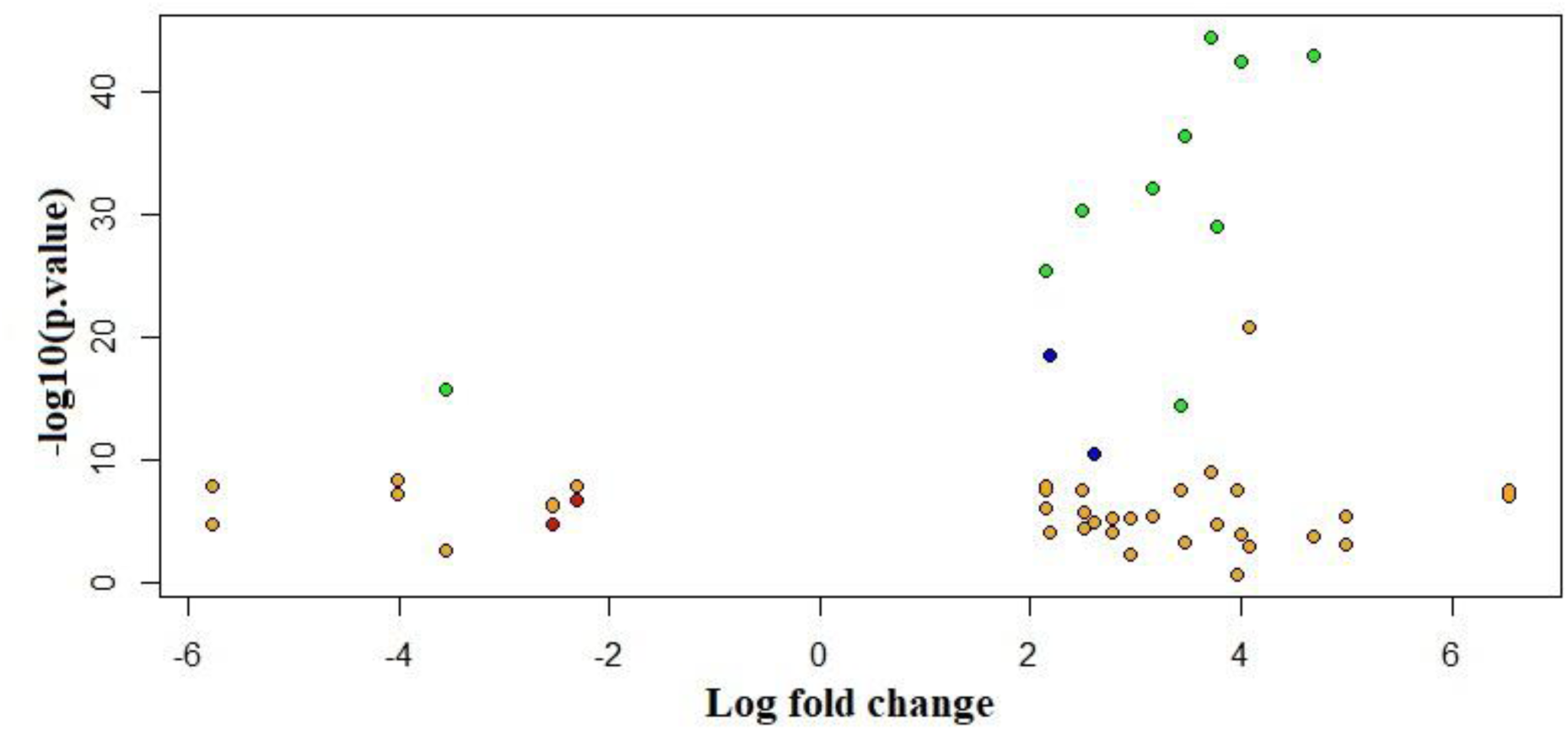
Volcano plot of final 49 significant stage-specific differenetially expressed genes. Stage 1 genes, red; Stage 2, blue; Stage 3, green; and Stage 4, orange. The genes are seen away from the origin, indicating significance and effect size.

**Table 6.** Number of genes in each step of the significance analysis. Differential expression is defined with respect to a threshold |logFC| = 2. Significance analysis proceeds first by significance (i.e, p-value) with respect to control, followed by p-value in each possible pairwise contrast between the different stages. Exclusive DE genes refer to genes differentially expressed in only one of the four stages (corresponding to the bit strings ‘1000’, ‘0100’, ‘0010’ and ‘0001’).

| Filtering criteria | STAGE 1 | STAGE 2 | STAGE 3 | STAGE 4 | Total |
| --- | --- | --- | --- | --- | --- |
| Exclusive DE genes | 40 | 75 | 223 | 481 | 819 |
| DE genes | 122 | 407 | 844 | 1082 | 2455 |
| Adj.p-value w.r.to control | 120 | 406 | 839 | 293 | 1658 |
| p-value 1 x 2 | 26 | 187 | - | - | 213 |
| p-value 1 x 3 | 19 | - | 670 | - | 689 |
| p-value 1 x 4 | 2 | - | - | 88 | 90 |
| p-value 2 x 3 | - | 13 | 70 | - | 83 |
| p-value 2 x 4 | - | 2 | - | 46 | 48 |
| p-value 3 x 4 | - | - | 10 | 35 | 45 |
| Final genes | 2 | 2 | 10 | 35 | 45 |

**Table 7.**
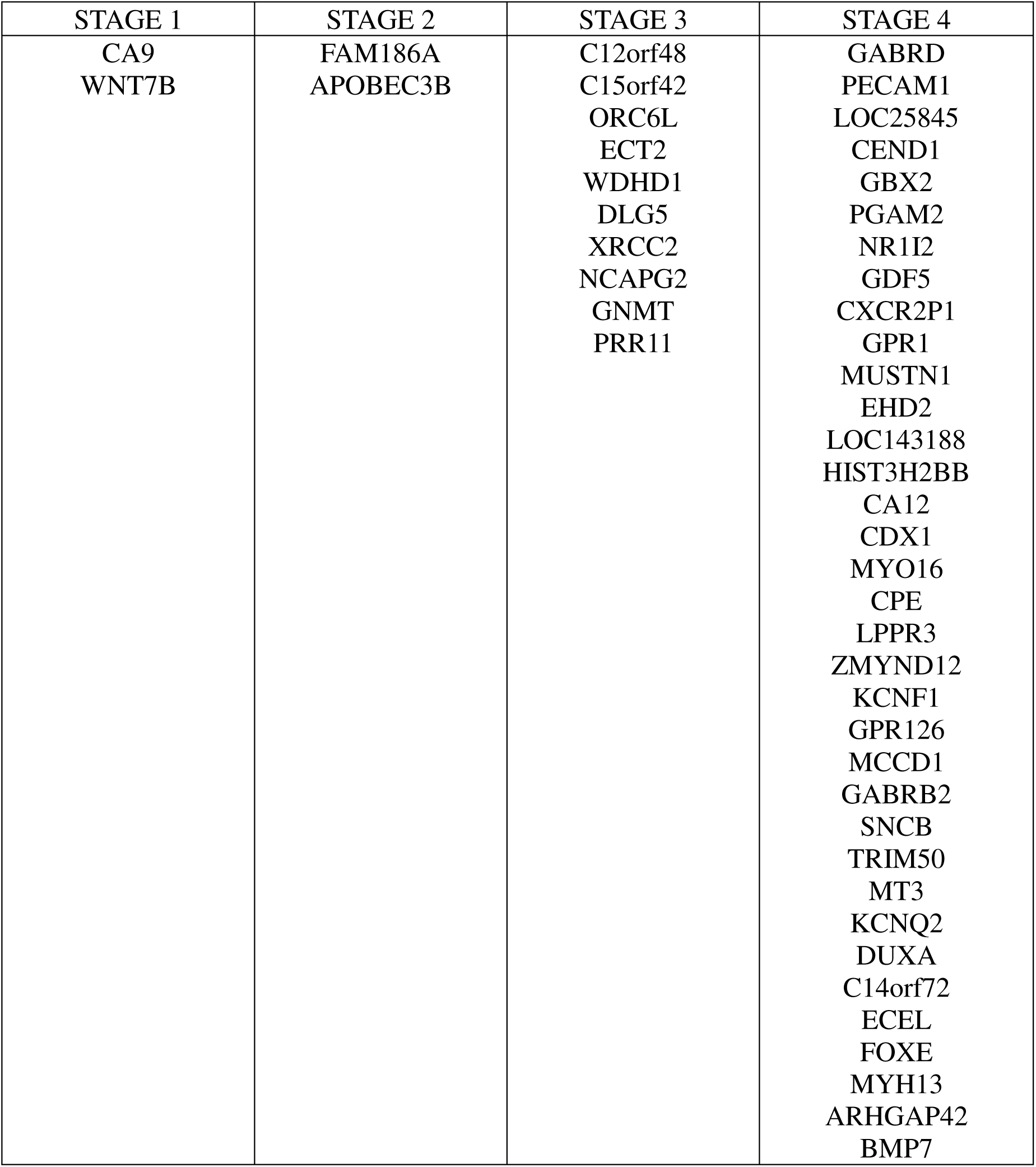
Final set of highlighted genes in each stage. The genes in each stage are ordered by increasing adjusted p-values of the linear modelling analysis.

In view of the limited sample size for stage-IV and consequent low power for rejecting false-positives, we stipulated that each stage-IV specific gene would display a smooth increasing or decreasing expression trend through cancer progression culminating in maximum differential expression in stage-IV. On this basis, we pruned the 35 stage-IV specific genes to just ten topranked by significance in the linear modelling.

A heatmap of the lfc expression of stage-specific genes across different stages was visualized (Fig. 10A) and revealed systematic variation in expression relative to control on a gradient from blue (downregulated) to red (overexpressed). The map was clustered on the basis of differential expression (i.e, |lfc|) both across stages and across features (i.e, genes) (Fig. 10B). Stage I genes clustered together, stage II genes co-clustered with NCAPG2 and DLG5 from stage-III, all the other stage-III genes clustered together, while the stage-IV genes formed two separate clusters. It was interesting to note that GABRD emerged as an outgroup to all the clusters, demonstrating its uniqueness.

**Figure 10.**
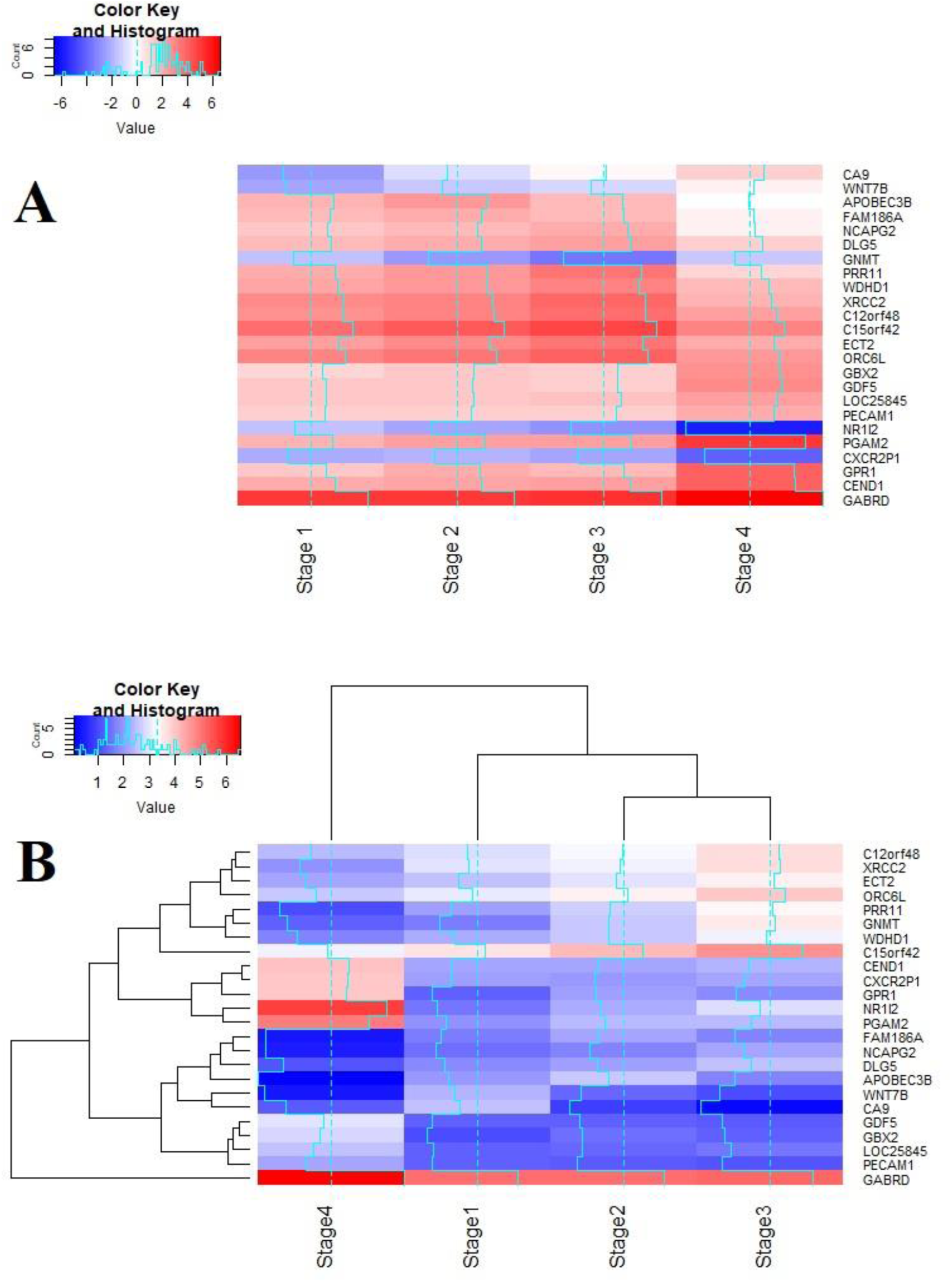
Heatmaps of final stage-specific 24 genes. **A,** heatmap generated from the lfc values of all the stage-specific genes (arranged stagewise). Log fold changes upto sixfold are seen, indicating 64 times differential expression with respect to control. **B,** Clustered representation of the stagewise gene expression based on differential expression (corresponding to |lfc|).

## DISCUSSION

When differentially expressed genes are identified in a two-class cancer vs control manner, the information about stage-specificity of differential expression is lost. By applying our protocol, this information is recovered and available for dissection.

To identify the biological processes specific to each stage, we used the genes with maximal |logFC| in each stage and performed a stagewise gene set enrichment analysis on two ontologies, the GO and KEGG pathways. Salient results with respect to KEGG pathways are presented below (Table 8) and the complete KEGG and GO results are available in Supplementary Tables S1 and S2, respectively. In stage I, we found the significant enrichment of cell-cycle signaling pathways (Hippo, Wnt, HIF-1), and viral infection-related pathways (cytokine-cytokine receptor interaction, human papillomavirus infection, HTLV-I infection). In stage II, key signalling pathways (Ras, MAPK) were aberrant. Two liver-specific pathways, alcoholism and cytochrome P450 mediated metabolism of xenobiotics were enriched, as well as standard cancer pathways of bladder, brain, stomach, and skin that might involve generic genetic alterations necessary for cancer cell growth. In stage III, we noticed the significant enrichment of Metabolic pathways that summarize cellular metabolism. This might indicate the metabolic shift needed by the cancer to grow and invade neighboring tissues. Other salient significantly enriched pathways pertained to increased cell cycle progression, DNA replication, chemical carcinogenesis, p53 signaling pathway and cellular senescence, all hallmark processes critical to cancer progression. Stage IV gene set was significantly enriched for bile-related processes (bile secretion, primary bile acid biosynthesis), and ABC transporters (possibly conferring a drug-resistant advanced cancer phenotype). A signaling pathway related to diabetic complications was enriched as well, indicating the role of co-morbidities in driving liver cancer progression. The enrichment analysis of the top 100 genes of the linear model is included in the Supplementary Table S3.

**Table 8.** Gene set enrichment analysis. Stage-specific gene sets (all the differentially expressed genes, corresponding to row ‘DE genes’ in Table 6) were analyzed for significant enrichment with respect to KEGG Pathways. Significance was based on p-value <0.05.

| Stage | Enriched pathways | p-value |
| --- | --- | --- |
| Stage 1 | Hippo signalling pathway | 3.276e-03 |
|  | Cytokine-cytokine receptor interaction | 1.218e-02 |
|  | Wnt signalling pathway | 1.528e-02 |
|  | Human papillomavirus infection | 1.763e-02 |
|  | HTLV-I infection | 2.552e-02 |
|  | HIF-1 signalling pathway | 2.787e-02 |
| Stage 2 | Bladder cancer | 4.643e-03 |
|  | Ras signalling pathway | 5.264e-03 |
|  | Pathways in cancer | 6.211e-03 |
|  | Glioma | 6.457e-03 |
|  | Alcoholism | 1.027e-02 |
|  | Gastric cancer | 1.210e-02 |
|  | MAPK signalling pathway | 2.526e-02 |
|  | Melanoma | 3.183e-02 |
|  | Metabolism of xenobiotics by cytochrome P450 | 3.472e-02 |
| Stage 3 | Cell cycle | 2.881e-18 |
|  | DNA replication | 6.526e-11 |
|  | Chemical carcinogenesis | 1.233e-06 |
|  | Metabolic pathways | 1.204e-03 |
|  | Cellular senescence | 7.203e-03 |
|  | p53 signalling pathway | 7.275e-03 |
| Stage 4 | Bile secretion | 2.479e-06 |
|  | ABC transporters | 7.146e-06 |
|  | Primary bile acid biosynthesis | 2.357e-03 |
|  | AGE-RAGE signalling pathway in diabetic complications | 3.024e-02 |

**Table 9.** Stage specific genes and parameters. The log-expressions (β’s) of each gene from our analysis are shown, along with adjusted p-values with respect to control and the linear model, and the inferred regulation status of the gene in LIHC. The stage-specificity of the genes are emphasized.

| GENE | $\beta_0$ | $\beta_1$ | $\beta_2$ | $\beta_3$ | $\beta_4$ | Adj.p.value<br>(from<br>contrasts<br>against<br>control) | Adj.p.value<br>(from linear<br>model) | Regulation<br>status<br>(Up/Down) |
| --- | --- | --- | --- | --- | --- | --- | --- | --- |
| STAGE 1 |  |  |  |  |  |  |  |  |
| CA9 | 0.1 | **-2.5** | -0.9 | 0.2 | 1.3 | 3.66e-05 | 3.44e-09 | Down |
| WNT7B | -1.8 | **-2.3** | -1.4 | -1.1 | 0.3 | 5.45e-07 | 2.07e-06 | Down |
| STAGE 2 |  |  |  |  |  |  |  |  |
| APOBEC3B | -0.9 | 2.0 | **2.6** | 1.7 | -0.6 | 1.12e-10 | 1.17e-09 | Up |
| FAM186A | -4.2 | 1.7 | **2.2** | 1.8 | 0.4 | 2.62e-18 | 1.50e-12 | Up |
| STAGE 3 |  |  |  |  |  |  |  |  |
| NCAPG2 | 1.5 | 1.5 | 1.8 | **2.1** | 0.4 | 6.03e-25 | 1.76e-24 | Up |
| DLG5 | 2.3 | 1.8 | 2.1 | **2.5** | 1.1 | 1.23e-29 | 2.90e-29 | Up |
| GNMT | 6.9 | -1.6 | -2.6 | **-3.6** | -1.3 | 9.69e-16 | 2.40e-15 | Down |
| PRR11 | -3.8 | 2.1 | 2.7 | **3.4** | 1.0 | 1.74e-14 | 4.65e-13 | Up |
| WDHD1 | -1.4 | 2.3 | 2.7 | **3.2** | 1.8 | 2.25e-31 | 8.87e-31 | Up |
| XRCC2 | -3.8 | 2.9 | 3.2 | **3.8** | 1.9 | 2.29e-28 | 7.06e-28 | Up |
| C12orf48 | -1.9 | 2.8 | 3.2 | **3.7** | 2.4 | 5.88e-43 | 1.19e-43 | Up |
| C15orf42 | -3.5 | 3.7 | 4.2 | **4.7** | 3.2 | 1.52e-41 | 1.17e-42 | Up |
| ECT2 | 0.2 | 2.5 | 3.0 | **3.5** | 2.2 | 2.62e-35 | 4.29e-35 | Up |
| ORC6L | -2.4 | 3.1 | 3.5 | **4.0** | 2.6 | 4.19e-41 | 5.55e-42 | Up |
| STAGE 4 |  |  |  |  |  |  |  |  |
| GABRD | -3.8 | 5.1 | 5.1 | 5.2 | **6.5** | 3.15e-17 | 5.53e-78 | Up |
| PECAM1 | 3.8 | 1.3 | 1.2 | 1.2 | **2.1** | 4.69e-07 | 7.64e-24 | Up |
| LOC25845 | 2.0 | 1.3 | 1.4 | 1.6 | **2.5** | 4.47e-06 | 1.29e-20 | Up |
| CEND1 | -5.0 | 2.2 | 2.2 | 2.4 | **4.1** | 4.42e-06 | 1.08e-17 | Up |
| GBX2 | -6.3 | 1.0 | 1.5 | 1.3 | **2.8** | 1.20e-05 | 2.59e-13 | Up |
| PGAM2 | -1.7 | 1.9 | 2.4 | 2.5 | **5.0** | 5.73e-05 | 1.72e-11 | Up |
| NR1I2 | 5.6 | -1.5 | -2.3 | -2.9 | **-5.8** | 8.41e-05 | 8.06e-11 | Down |
| GDF5 | -6.1 | 1.3 | 1.3 | 1.2 | **2.9** | 2.73e-05 | 8.22e-11 | Up |
| CXCR2P1 | 2.0 | -2 | -2 | -2.2 | **-4.0** | 4.52e-04 | 1.77e-10 | Down |
| GPR1 | -5.5 | 1.3 | 2.1 | 1.8 | **3.9** | 1.42e-04 | 4.33e-10 | Up |

The top ten linear model genes (Table 5) and all the stage-specific differentially expressed genes (Table 10) were analyzed with respect to the existing literature. Three C-type lectin domain proteins (CLEC4M, CLEC1B, CLEC4G) were detected in the top ten genes of linear modelling across stages. Interestingly, this identical cluster of three genes was detected as the most significantly downregulated liver cancer-specific genes in a qPCR study of an independent cohort of 65 tumor-normal matched cases (Ho et al., 2015). On screening the top 200 linear model genes against cancer driver genes in the Cancer top 200 Gene Census, only four genes were found, namely BUB1B, CDKN2A, EZH2, and RECQL4.

*Stage-I specific DEGs* (Fig. 11). CA9 is a member of carbonic anhydrases, which are a large family of zinc metalloenzymes that catalyse the reversible hydration of carbon dioxide. Its expression in clear cell Renal carcinoma, but not in functional kidney cells has gained attention for its use as a pre-operative biomarker (Li et al., 2017). The WNT7B protein is part of the Wnt family, a family of secreted signalling proteins. Elevated WNT7B in pancreatic adenocarcinoma has been found to mediate anchorage independent growth (Arensman et al., 2014). Surprisingly, both CA9 and WNT7B are downregulated in LIHC, most so in stage-I, contrary to their role in other cancers. A concrete interpretation of the role of these genes in LIHC awaits appropriately designed experimental studies.

**Figure 11.**
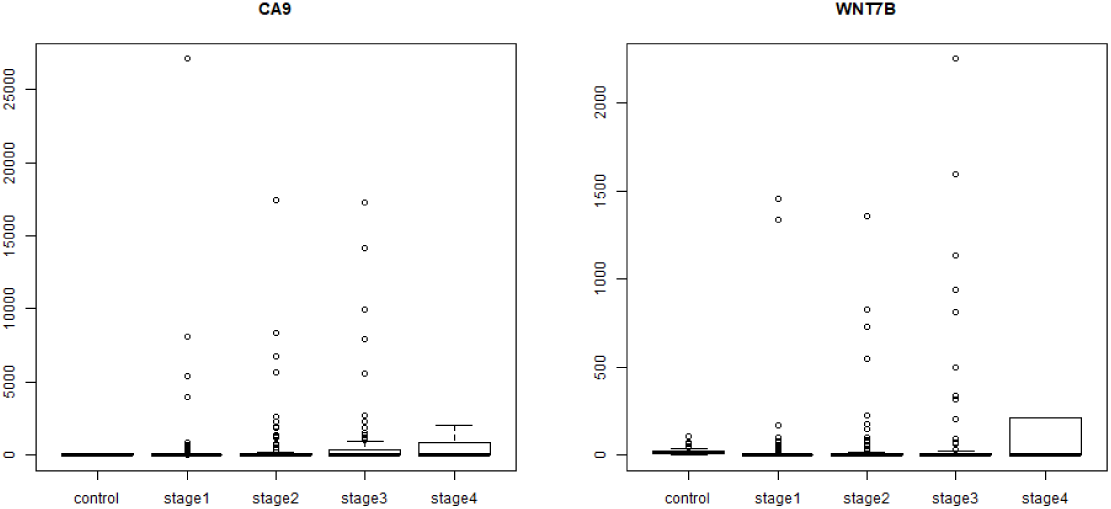
Boxplot of stage-I specific genes. It is seen that CA9 and WNT7B are both maximally downregulated in stage-I.

It is pertinent to ask the following question here: which genes are essential for the initiation of LIHC? Clearly these genes would be differentially expressed in stage I relative to control. All significantly differentially expressed genes with maximal |lfc| in stage-I would be the best candidates for genes involved in the initiation of LIHC. These 122 genes are provided in the Supplementary File S3.

*Stage-II specific DEGs* (Fig. 12). APOBEC3B, a DNA cytidine deaminase, is a known cancer driver gene in the Cancer Gene Census, but there are no literature reports of its stage-specificity in any cancer. It is known to accountfor half the mutational load in breast carcinoma, and its target sequence was found to be highly mutated in Bladder, lung, cervix, neck, and head cancers as well (Burns et al., 2013). Here APOBEC3B is upregulated possibly conferring a gain-of-function comparable to that achieved by a mutation mechanism. FAM186A polymorphisms have been reported in GWAS and SNP studies on colorectal cancer patients and shown to have a significant odds ratio in risk heritability (Timofeeva et al., 2015).

**Figure 12.**
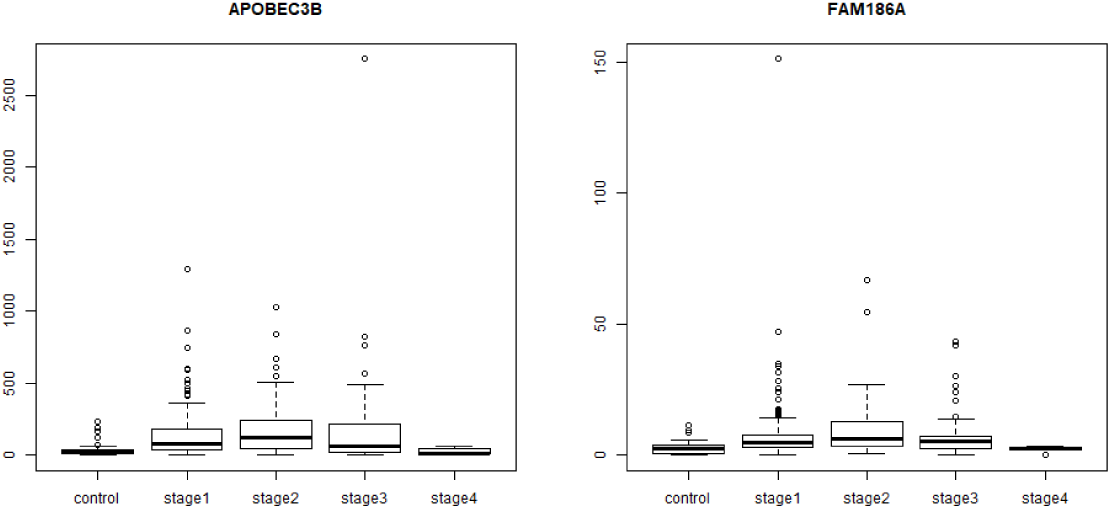
Boxplot of stage-II specific genes. It is seen that both APOBEC3B and FAM186A expression are maximum in stage-II, following an inverted U-shape.

*Stage-III specific DEGs* (Fig. 13). C12orf48, also known as PARI, participates in the homologous recombination pathway of DNA repair, and its overexpression has been reported in pancreatic cancer(O’Connor et al., 2013). Further PARI was recently identified as a transcriptional target of FOXM1 (Zhang et al., 2018), which is a well-validated upregulated gene in LIHC (Ho et al., 2015). DLG5 is a cell polarity gene and its downregulation has been implicated in the malignancy of breast (Liu et al., 2017), prostate (Tomiyama et al., 2015) and bladder cancers (Zhou et al., 2015). It has been recently found that lower DLG5 expression is correlated with advanced stages of HCC and essential for invadopodium formation, an event critical to cancer metastasis (Ke et al., 2017). It is surprising that our study has identified a stage-III specific upregulation in DLG5. Interestingly, evidence is emerging to lend support to our finding that DLG5 might be tumor-promoting. In a very recent review, Saito et al. (2018) reinterpreted published results on cell polarity and cancer, and advanced an alternative perspective on the role of polarity regulators in cancer biology. They argued that both cellular and subcellular polarity would be regulated by DLG5 and related polarity proteins. Subcellular polarity might improve the cellular fitness for proliferation and stemness, thereby causing tumor promotion. Hence cell polarity regulation is anti-tumorigenic and subcellular polarity regulation is pro-tumorigenic, and our analysis has uncovered the pro-tumorigenic upregulated activity of DLG5.. ECT2 encodes a guanine nucleotide exchange factor that remains elevated during the G2 and M phase in cellular mitosis. ECT2 is found to be upregulated in lung adenocarcinoma and lung squamous cell carcinoma (Zhou et al., 2017), as well as in invasive breast cancer (Wang et al., 2017). NCAPG2 is a component of the condensing II complex and involved in chromosome segregation during mitosis. NCAPG2 level were found to be increased in non-small cell lung cancer, and its over-expression was found to be correlated with lymph node metastasis, thus enabling the use of NCAPG2 as a poor prognostic biomarker in lung adenocarcinoma (Zhan et al., 2017). GNMT is a methyltransferase that catalyses conversion of S-adenosine methionine to sadenosyl cysteine. In the absence of GNMT, S-adenosine methionine causes hypermethylation of DNA, which represses GNMT levels and is found in HCC samples (Huidobro et al., 2013). This is an epigenetic mechanism for loss of function of tumor suppressors and our study here confirmed the downregulation of GNMT expression. PRR11 is found to be over-expressed in lungs, and its silencing using siRNA resulted in cell cycle arrest and apoptotic cell death, followed by decreased cell growth and viability (Zhao 2015). A similar knock out experiment of PRR11 in hilar cholangiocarcinoma cell lines resulted in decreased cellular proliferation, migration, and tumor growth (Chen et al., 2015). WDHD1 is a key post-transcriptional regulator of centromeric, and consequently genomic, integrity (Hsieh et al., 2011) and its overexpression has been identified as biomarker of acute myeloid leukemia (Wermke et al., 2015), and lung and esophageal carcinomas (Sato et al., 2010). C15orf42 has been implicated in nasopharyngeal carcinoma (An et al., 2015). ORC6L overexpression has been identified as a prognostic biomarker of colorectal cancer possibly by enhancing chromosomal instability (Xi et al., 2008). XRCC2 was found to increase locally advanced rectal cancer radioresistance by repairing DNA double-strand breaks and preventing cancer cell apoptosis (Qin et al 2015).

**Figure 13.**
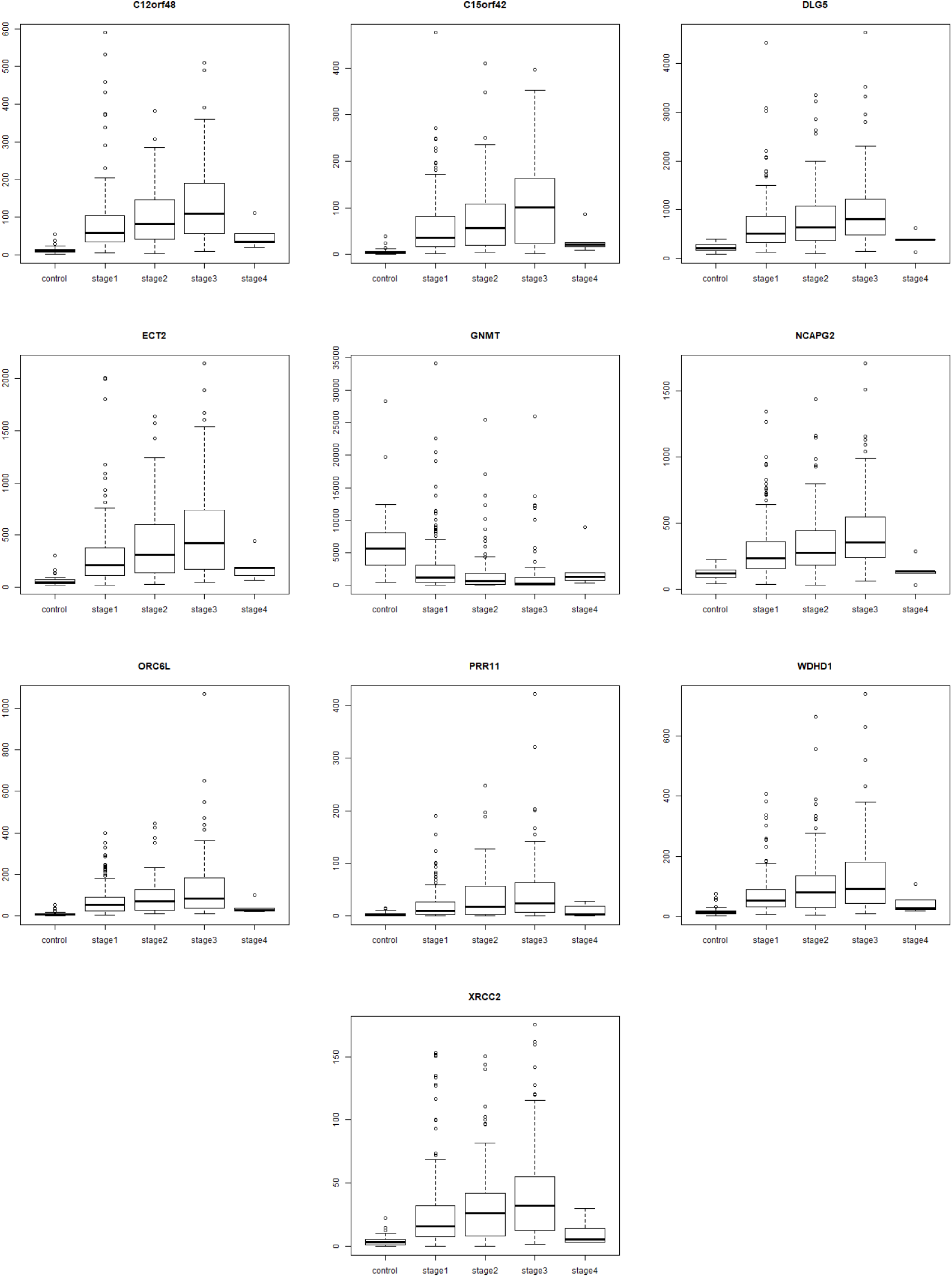
Boxplot of stage-III specific genes. Except for GNMT, the expression of all other genes show a in stage-III, following an inverted U-shape. The expression trend is reversed for the downregulated GNMT, following a U-shape.

*Stage-IV specific DEGs* (Fig. 14). GABRD, which is the top gene in the linear model as well, encodes for the delta subunit of the gamma-amino butyric acid receptor. The GABA receptor family was found to be frequently downregulated in cancers, except for GABRD, which was found to be up-regulated. Gross et al. (2015) proposed that the GABA receptor gene family might play a role in the proliferation independent differentiation of cancer cells. GBX2 is part of the GBX gene family, which are homeobox containing DNA binding transcription factors. GBX2 is overexpressed in prostate cancer and studies show that expression of GBX2 is required for malignant growth of human prostate cancer (Gao et al., 1998). PECAM1 overexpression has been linked to peritoneal recurrence of stage II/III gastric cancer patients (Terashima et al., 2017). CEND1 has been identified as a cell-cycle protein (Tsioras et al., 2013). PGAM2 is a glycolytic enzyme whose upregulation is essential for tumor cell proliferation (Xu et al., 2014). NR1I2 downregulation has been used in constructing a prognostic 9-genes expression signature of gastric cancer (Wang et al., 2017). GDF5 has been shown to be a downstream target of the TGF-beta signaling pathway (Margheri et al., 2012), stimulating angiogenesis required for the growth and spread of the cancer. GPR1 has been reported to be involved in promoting cutaneous squamous cell carcinoma migration (Farsam et al., 2016). Two more stage-IV specific genes, namely CXCR2P1, which is a C-X-C motif chemokine receptor 2 pseudogene 1, and LOC25845, are undocumented in the literature in the context of LIHC, other cancers or any other condition.

**Figure 14.**
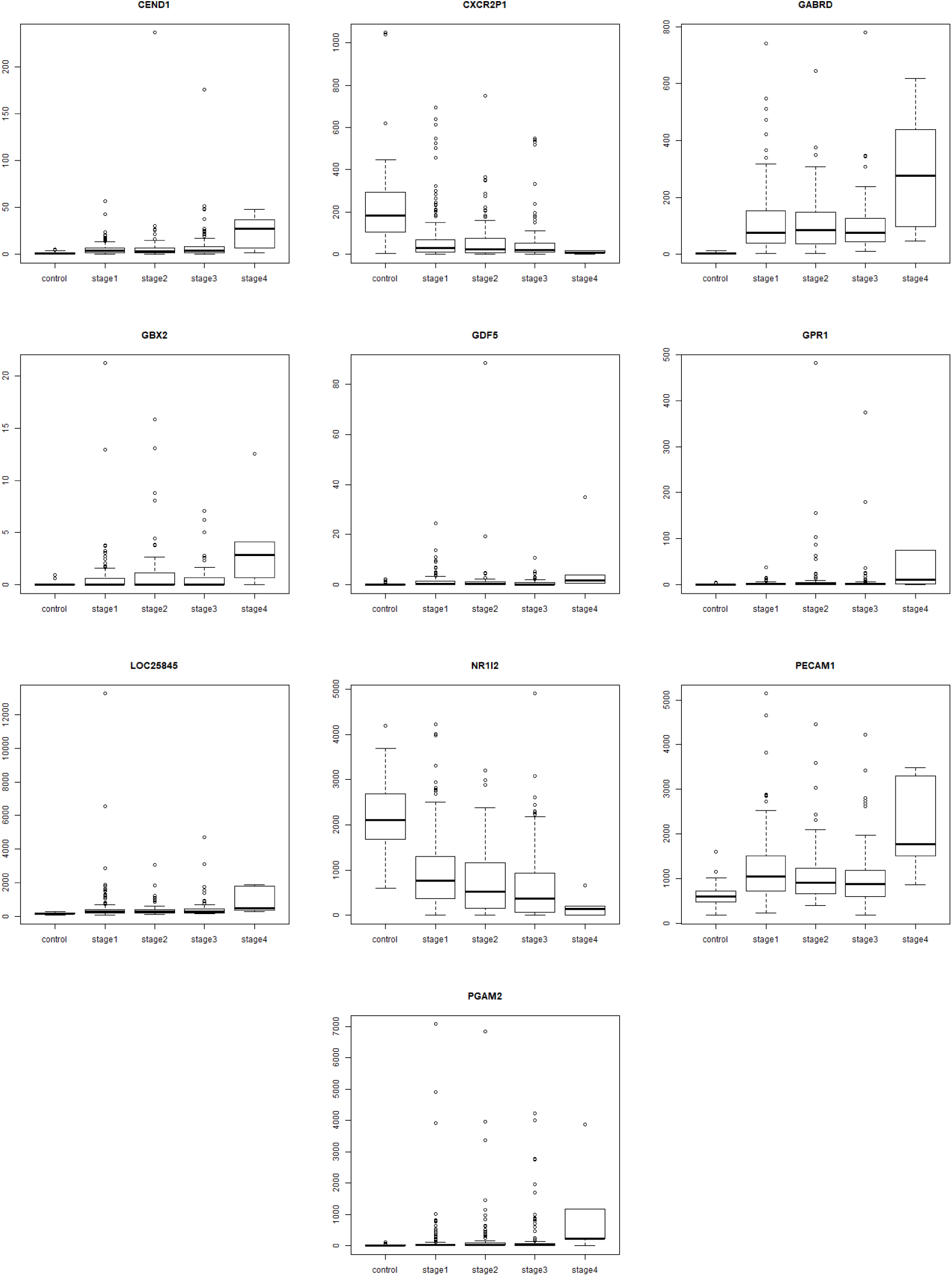
Boxplot of top 10 stage 4 specific genes. All genes, except NR1I2 and CXCR2P1, show a smooth increasing trend of expression reaching its maximum in stage-IV. For NR1I2 and CXC2RP1, the trend is reversed, with a smooth decreasing expression reaching its minimum in stage-IV.

## CONCLUSION

We have developed an original protocol for the stagewise dissection of the LIHC transcriptome. We were able to successfully fit a linear model across cancer stages and detected genes with a strong linear expression trend in the cancer phenotype. These genes were found to effectively separate the control and cancer samples. We were able to assign 2455 differentially expressed genes into one of four stages and visualized their stage specific expression using boxplots. Using a multi-layered approach, we were able to assess the significance of each stage-specific DEG and narrowed down to a handful of candidate significant stage-specific DEG’s. Our analysis yielded two stage-I specific genes (CA9, WNT7B), two stage-II specific genes (APOBEC3B, FAM186A), ten stage-III specific genes and ten stage-IV specific genes. Though all these genes except APOBEC3B are novel, a literature search indicated that most of the genes have a cancer connection (albeit not with LIHC). Experimental validation would be useful to translate these results into a panel of biomarkers for clinical use and rational drug development. It is straightforward to extend our computational methodology to the stage-based analysis of other cancers to obtain a fuller view of disease initiation, progression, and metastasis.

## ACKNOWLEDGMENTS

We would like to thank SASTRA deemed University, for infrastructure and computing support.

## SUPPLEMENTARY INFORMATION

All data and results are provided in the supplementary information (10.6084/m9.figshare.6455024).

